## Supplemental figures for "Evolutionary and functional analyses reveal a role for the RHIM in tuning RIPK3 activity across vertebrates"

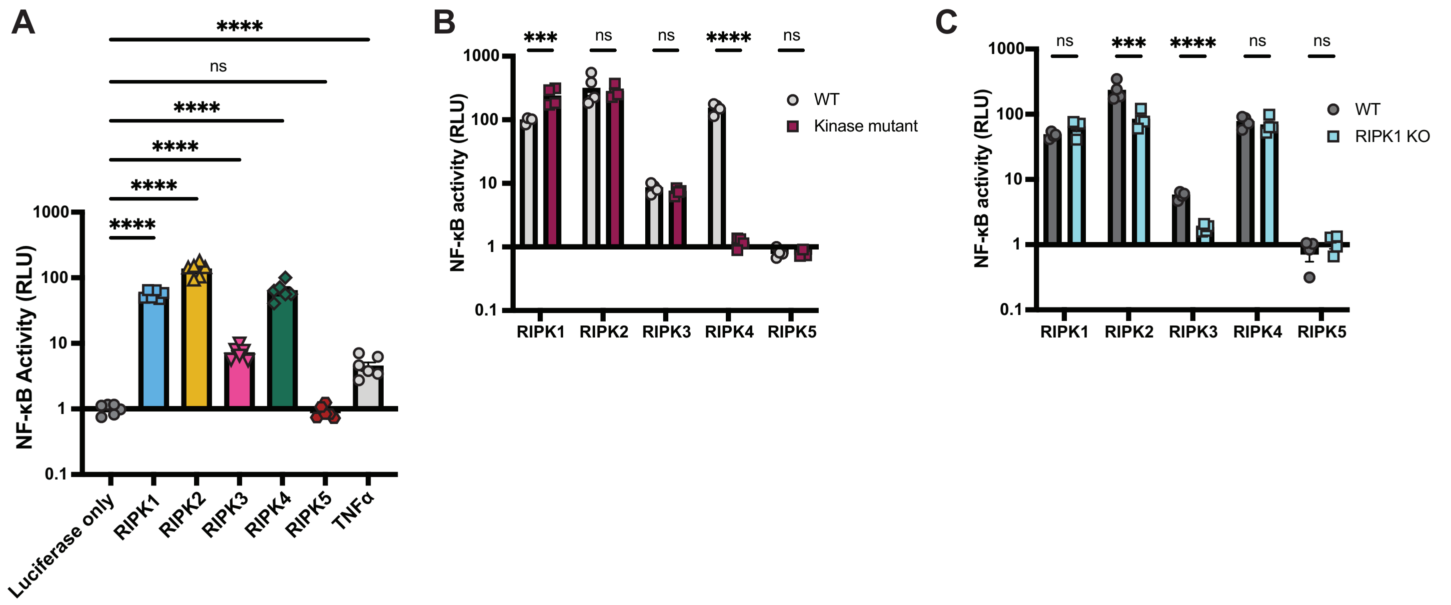


**Supplementary figure 1.** Human RIPK1-4 activate NF-κB. Human V5-RIPK1-5 proteins were transfected into WT (A-C) or RIPK1 KO (C) HEK293T cells, along with NF-κB firefly luciferase and control renilla luciferase reporter plasmids (see Materials and Methods), and NF-κB activity was measured at 18h post-transfection. (A) Human RIPK1-5 activation of NF-κB in WT 293T cells. (B) Activation of NF-κB by catalytically active versus kinase-mutant RIPK1-5 in WT HEK293T cells. (C) Activation of NF-κB by RIPK1-5 in WT versus RIPK1 KO HEK293T cells) Data are representative of 3 independent experiments with n=3-6 replicates per group. Data were analyzed using one-way ANOVA with Dunnett's multiple comparisons test (A) or two-way ANOVA with Šidák's multiple comparisons test (B, C). ns = not significant, *** = p<0.001, **** = p<0.0001.

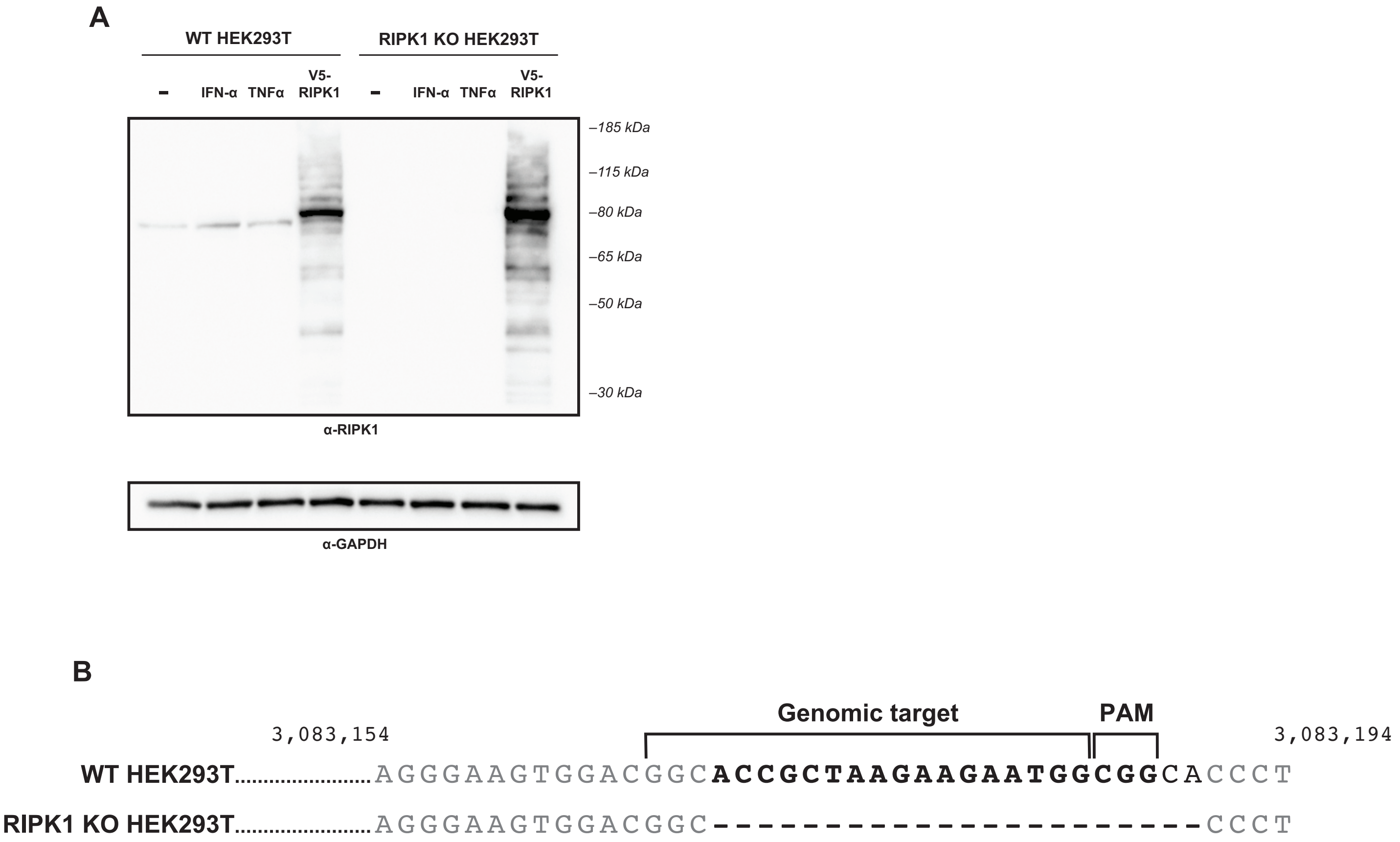


**Supplementary figure 2.** Generation of RIPK1 KO HEK293T cells. (A) WT and RIPK1 KO HEK293T cells were treated with 5000U IFN-$\alpha$ for 24 hours or 160 ng/mL TNF$\alpha$ for 6 hours and expression of RIPK1 protein was analyzed by western blot using the indicated antibodies. Untreated cells were used as a negative control. Cells were transfected with V5-RIPK1 as a positive control. (B) Genomic DNA was isolated from WT HEK293T (top) or RIPK1 KO HEK293T (bottom) cells and the target region of exon 5 of RIPK1 was amplified and analyzed by Sanger sequencing. Nucleotide numbers indicate genomic location.

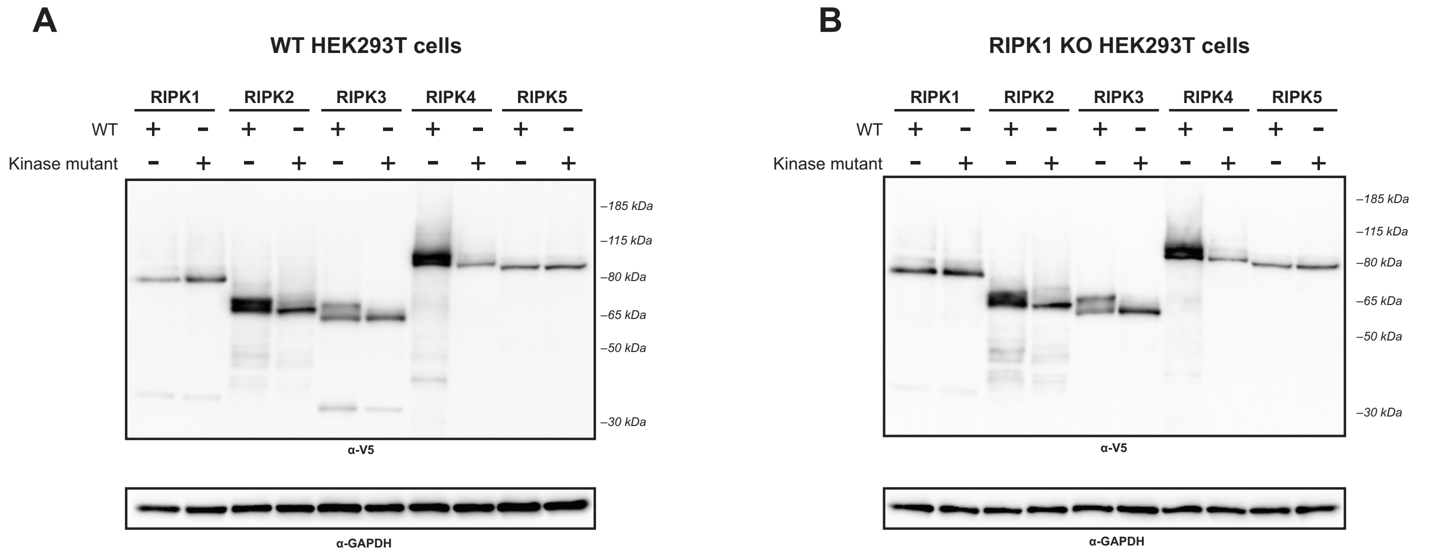


**Supplementary figure 3.** Expression of RIPK1-5 constructs. WT (A) and RIPK1 KO (B) HEK293T cells were transfected with WT or kinase mutant V5-RIPK1-5. Protein expression was analyzed at 18h post-transfection by western blot using the indicated antibodies.

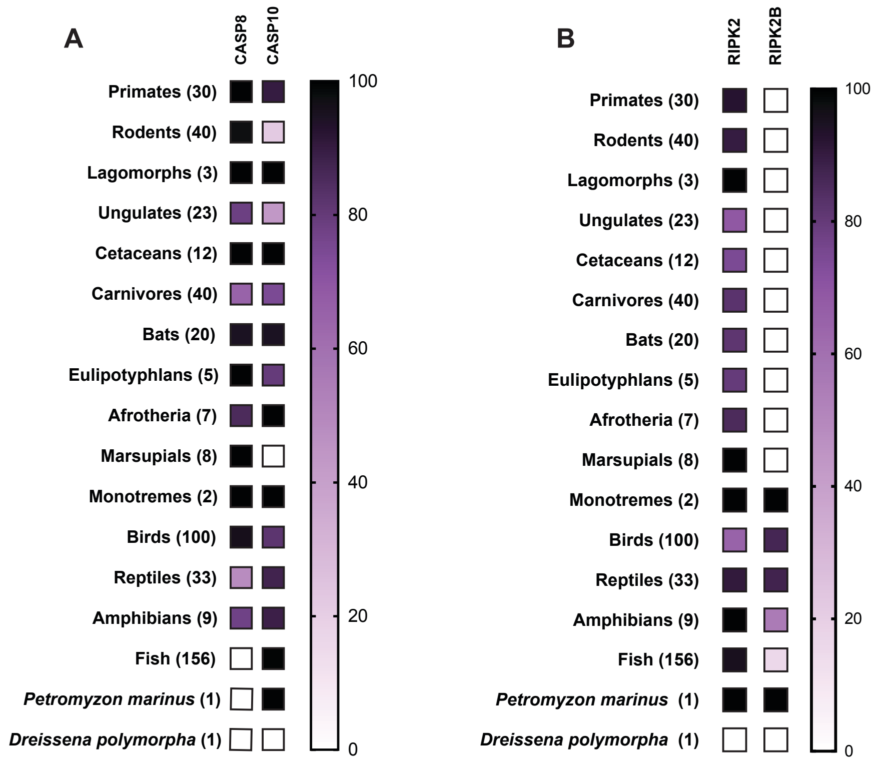


**Supplementary figure 4.** (A) Delineation between CASP8 and CASP10 as determined by phylogeny (see Materials and Methods). (B) Presence of RIPK2 and RIPK2B in vertebrates. Accession numbers for proteins analyzed and list of species can be found in Supplementary Files 3-4.

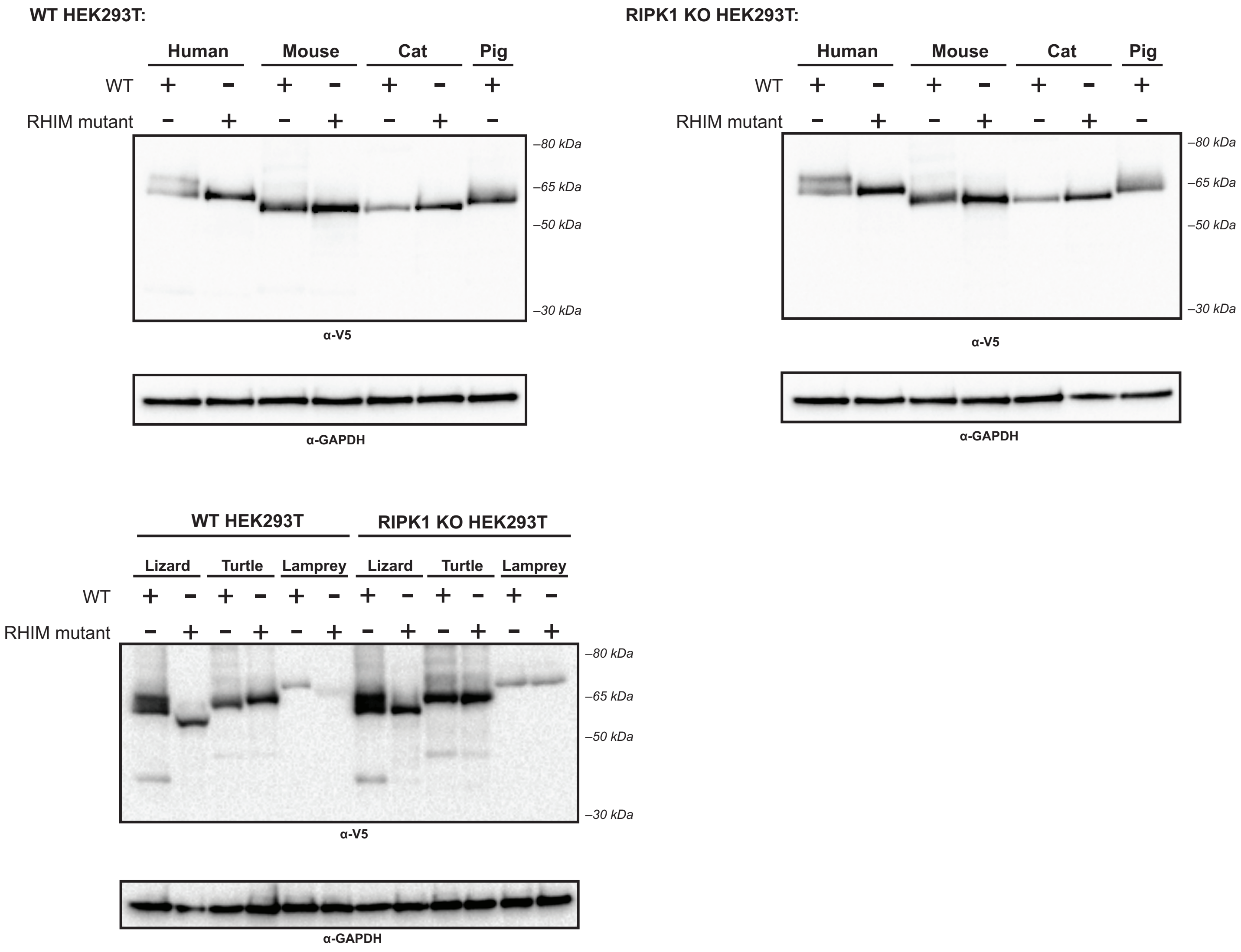


**Supplementary figure 5.** Expression of non-human RIPK3 proteins. WT or RIPK1 KO HEK293T cells were transfected with the indicated protein. Protein expression was analyzed at 18 hours post-transfection by western blot using the indicated antibodies.

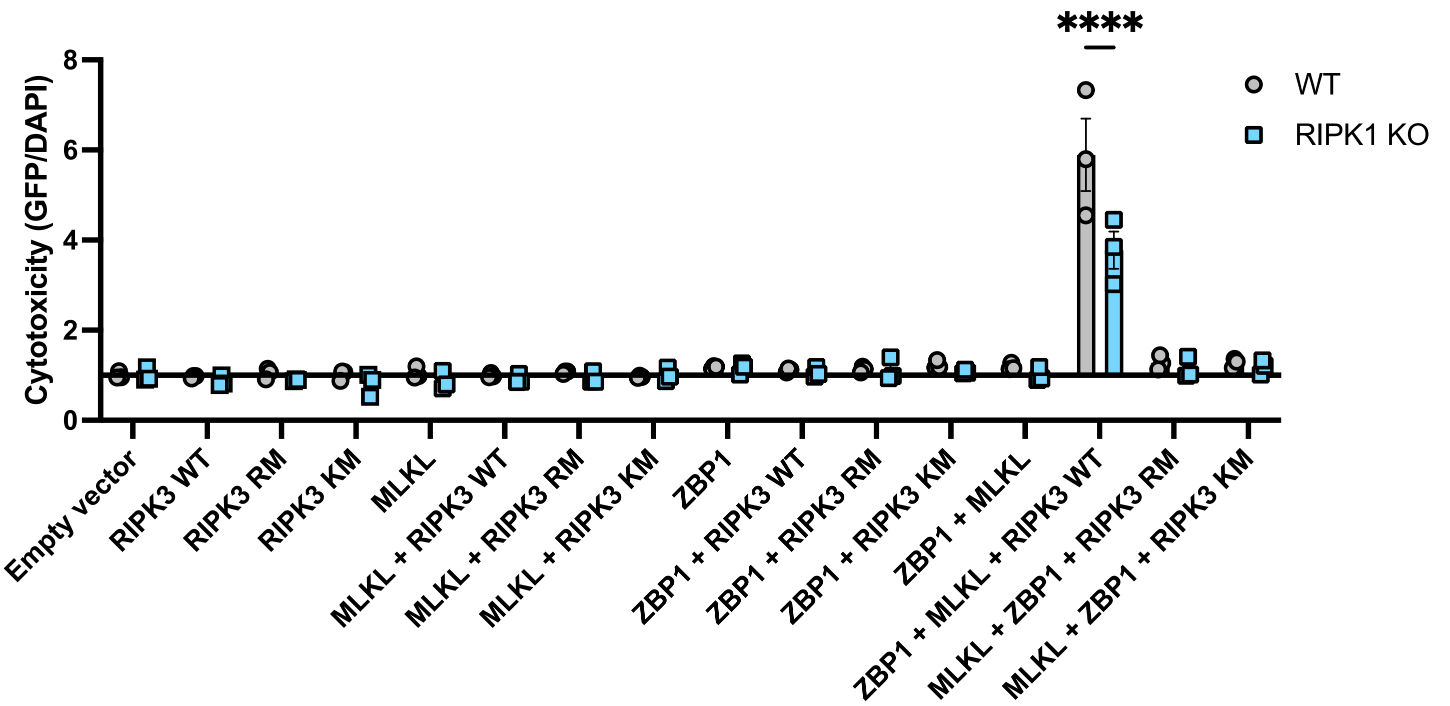


**Supplementary figure 6.** MLKL- and ZBP1-dependent cell death activation by RIPK3. WT and RIPK1 KO cells were transfected with the indicated plasmids and cell death was analyzed by ReadyProbes assay at 18h post-transfection. RM = RHIM mutant, KM = kinase mutant. Data indicative of 1-2 independent experiments with n=3 replicates per group. Data were analyzed by two-way ANOVA with Šidák's multiple comparisons test. **** = p<0.0001.

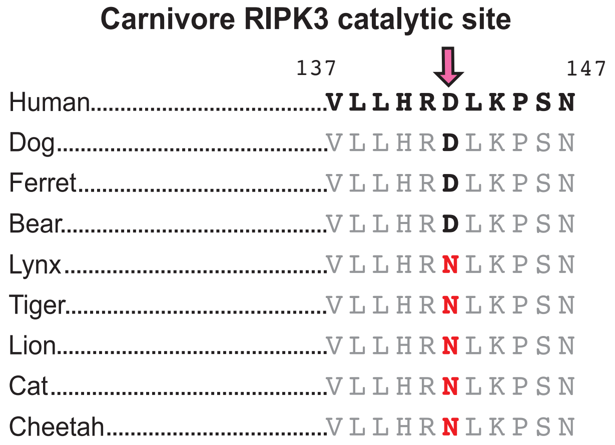


**Supplementary figure 7.** Loss of RIPK3 catalytic site in carnivores. RIPK3 sequences from carnivore species highlighting the catalytic site in the kinase domain. Residue numbers refer to the human sequence. Carnivore species include dog (*Canis lupus familiaris*), ferret (*Mustela putorius furo*), bear (*Ursus americanus*), lynx (*Lynx canadensis*), tiger (*Panthera tigris*), lion (*Panthera leo*), cat (*Felis catus*), cheetah (*Acinonyx jubatus*).

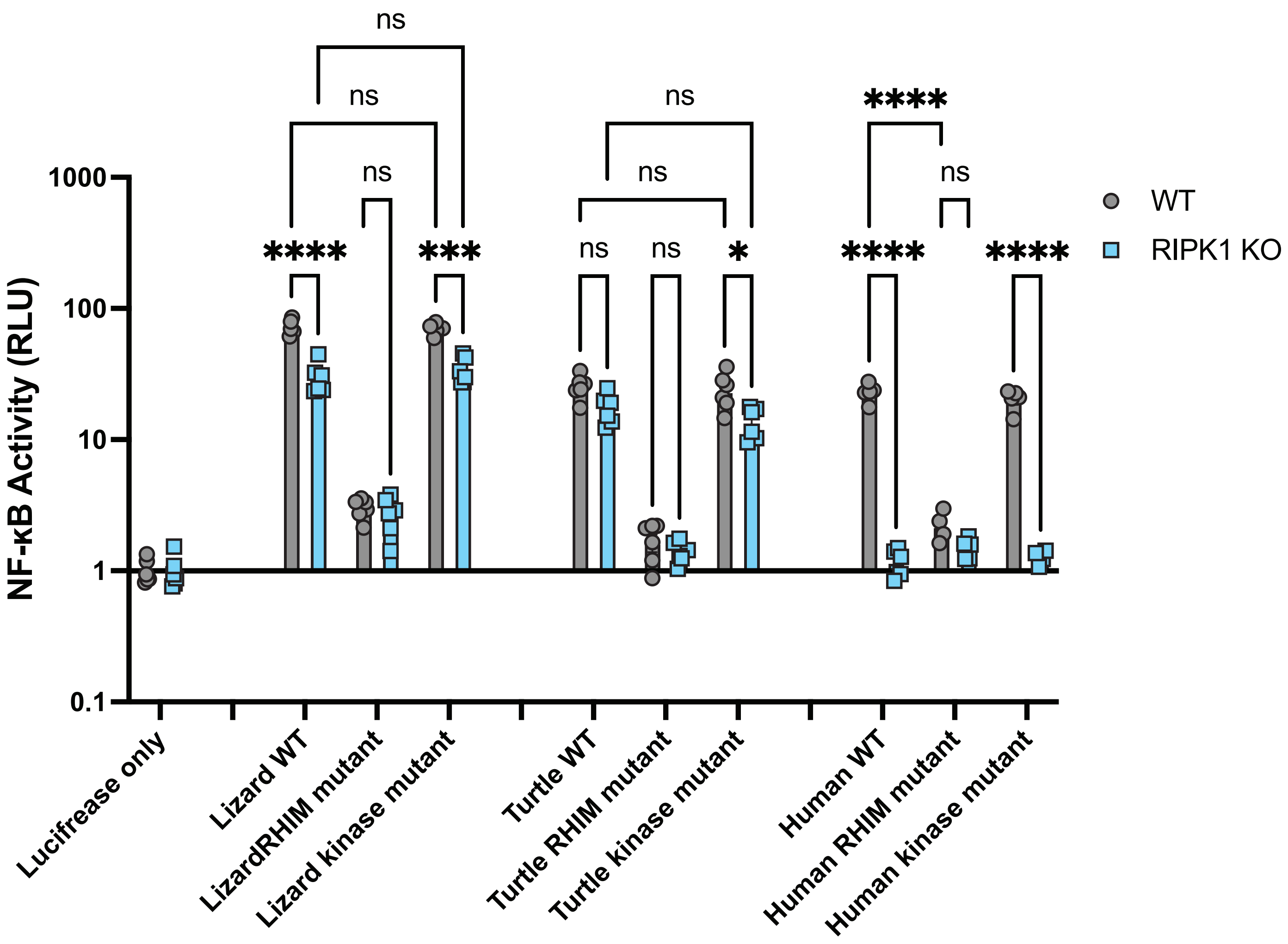


**Supplementary figure 8.** RIPK1-independent activation of reptile RIPK3 is independent of kinase activity. The indicated RIPK3 proteins were transfected into WT or RIPK1 KO HEK293T cells, along with NF-κB firefly luciferase and control renilla luciferase reporter plasmids (see Materials and Methods), and NF-κB activity was measured at 18h post-transfection. Data representative of two independent experiments with n=4-6 replicates per group. Data were analyzed using 2-way ANOVA with Tukey's multiple comparison's test. ns = not significant, * $p<0.05$, *** $p<0.001$, **** $p<0.0001$

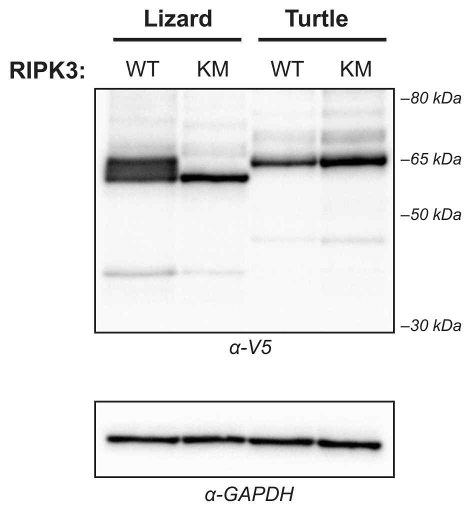


**Supplementary figure 9.** Expression of reptile RIPK3 kinase mutants. WT HEK293T cells were transfected with the indicated V5-RIPK3 protein. Protein expression was analyzed at 18 hours post-transfection by western blot using the indicated antibodies. WT = wild type, KM = kinase mutant.

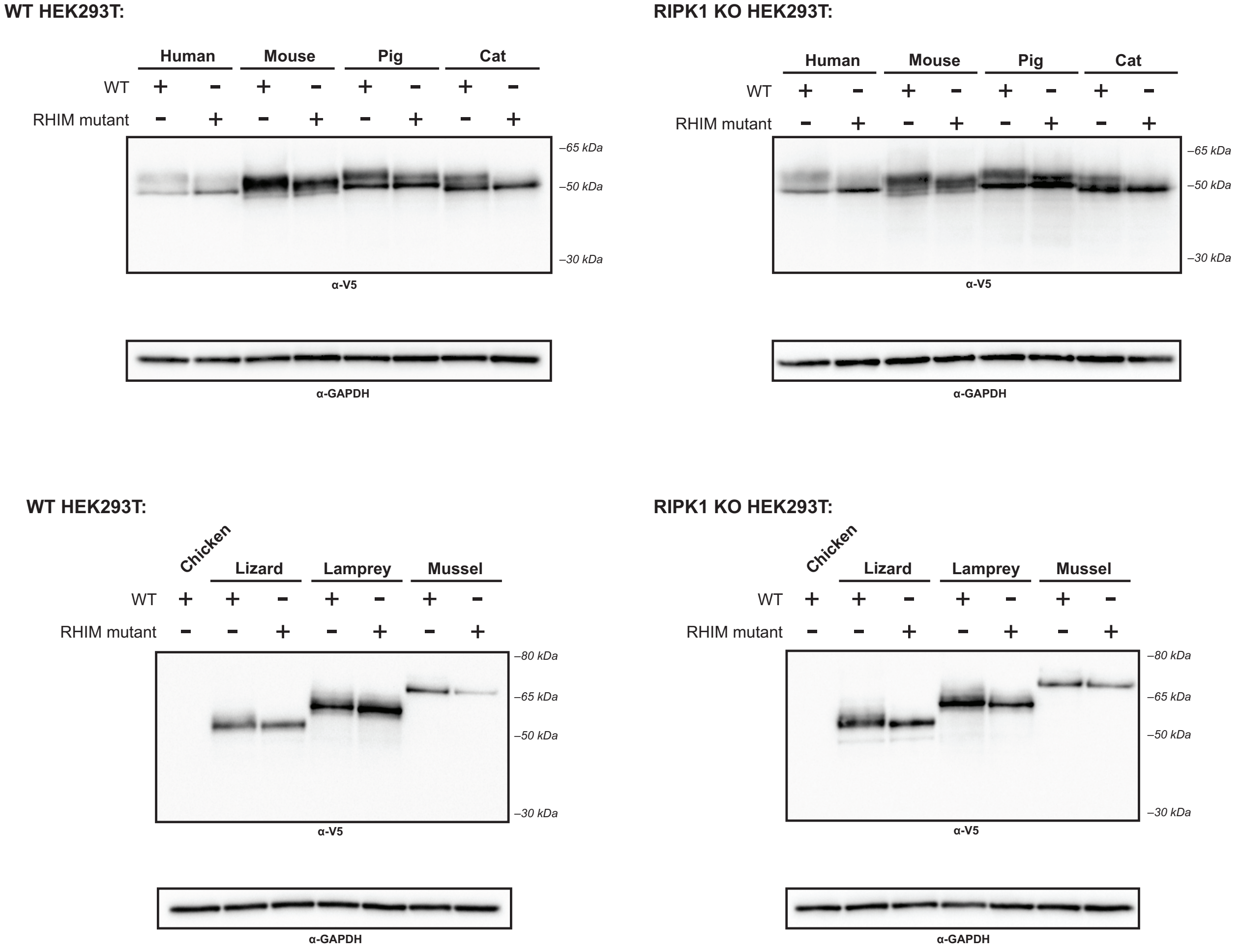


**Supplementary figure 10.** Expression of non-human RIPK1$^{CT}$ proteins. WT or RIPK1 KO HEK293T cells were transfected with the indicated protein. Protein expression was analyzed at 18 hours post-transfection by western blot using the indicated antibodies.

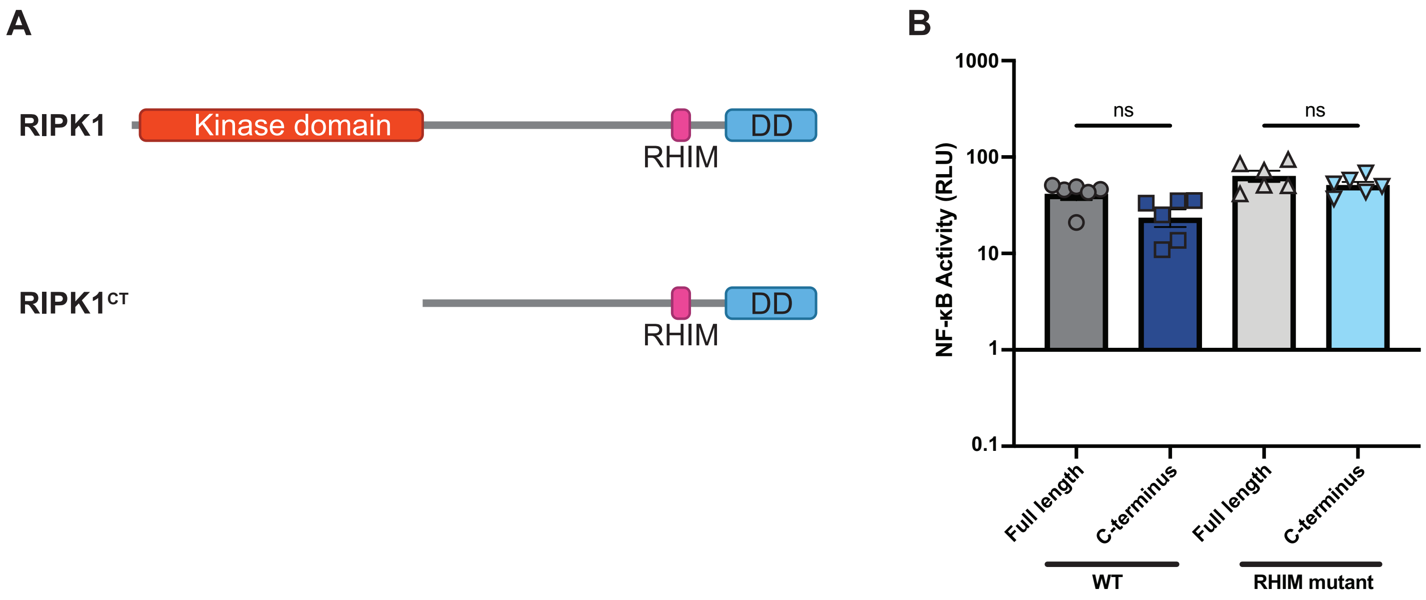


**Supplementary figure 11.** RIPK1$^{CT}$ activates NF-κB similar to full length RIPK1. (A) Schematic of RIPK1$^{CT}$ compared to full length RIPK1. (B)WT and RHIM mutant RIPK1 full length and RIPK1$^{CT}$ were transfected into HEK293T cells along with Dual-Glo plasmids (see Methods, Figure 5). NF-κB activation was analyzed at 18h post-transfection. Data were analyzed using one-way ANOVA with Tukey's multiple comparison's test. ns = not significant.

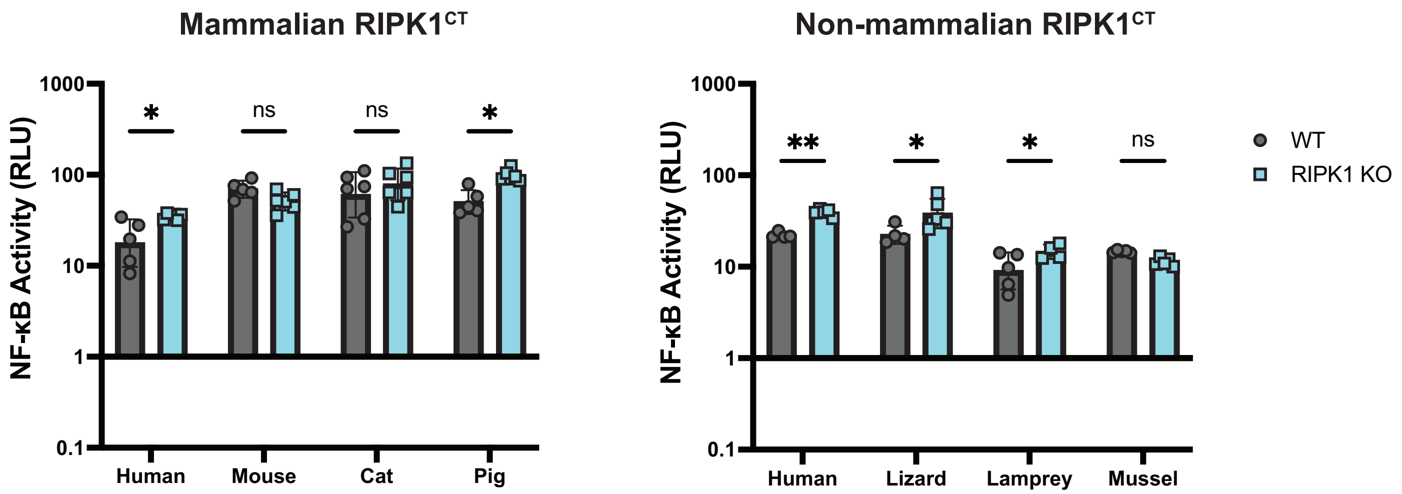


**Supplementary Figure 12**. Activation of NF-κB by diverse vertebrate RIPK1$^{CT}$ is independent of endogenous human RIPK1. WT and RIPK1 KO HEK293T cells were transfected with the indicated RIPK1$^{CT}$ along with Dual-Glo plasmids (see Methods).NF-κB activation was analyzed at 18h post transfection. Data representative of three independent experiments with 3-6 replicates per group. Data were analyzed using two-way ANOVA with Šidák's multiple comparisons test. ns = not significant, * = p<0.05, ** = p<0.01.

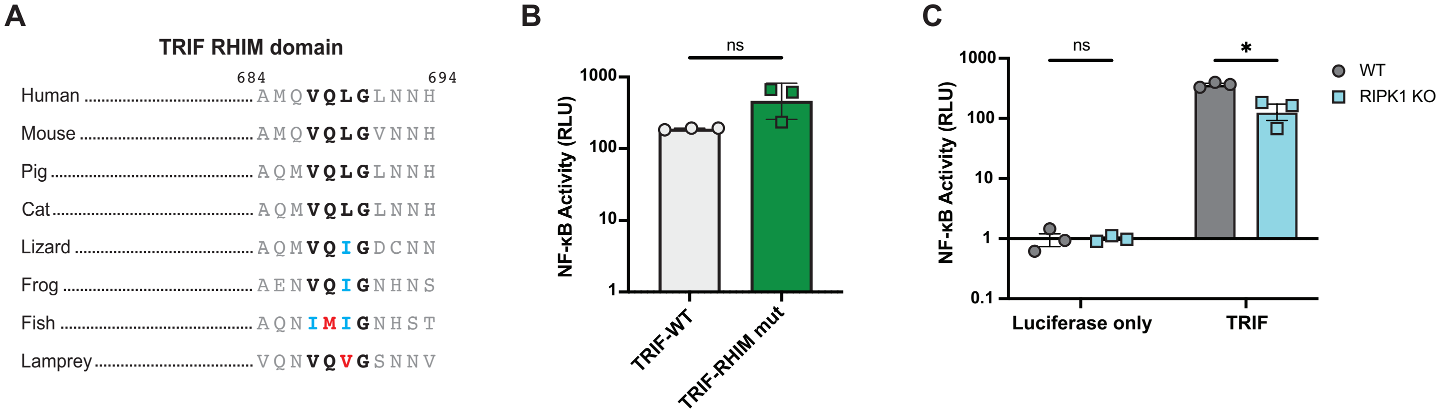


**Supplementary Figure 13.** Activation of NF-κB by TRIF. (A) TRIF sequences from vertebrate species highlighting the RHIM core tetrad. Residue numbers refer to the human sequence. Vertebrate species shown include mouse (*Mus musculus*), pig (*Sus scrofa*), cat (*Felis catus*), lizard (*Anolis carolinensis*), frog (*Xenopus laevis*), fish (*Danio rerio*), and lamprey (*Petromyzon marinus*. (B-C) WT (B-C) and RIPK1 KO (C) HEK293T cells were transfected with the indicated plasmids along with Dual-Glo plasmids (see Methods). NF-κB activation was analyzed at 18h post transfection. Data representative of three independent experiments with 3 replicates per group. Data were analyzed using a t-test (B) or a two-way ANOVA with Šidák's multiple comparisons test (C). ns = not significant, * = $p<0.05$.

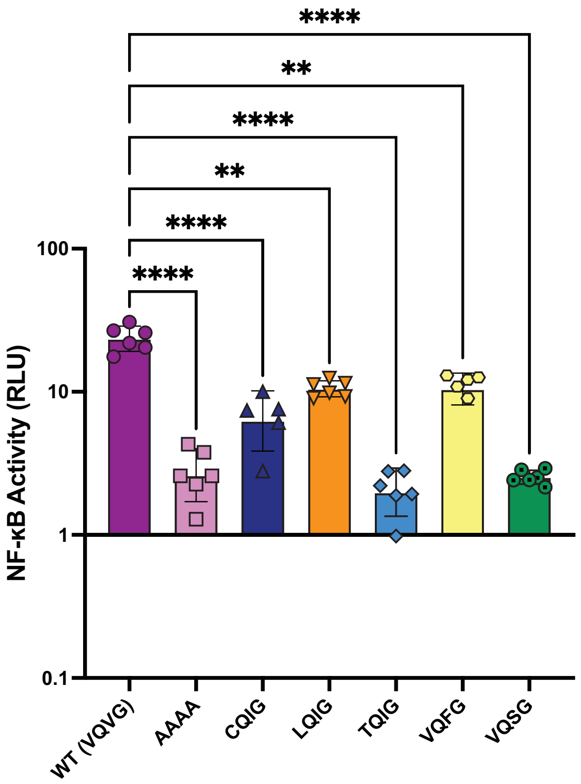


**Supplementary figure 14.** Activation of NF-κB by human RIPK3 RHIM variants. The indicated human RIPK3 RHIM tetrad variants were transfected into HEK293T HEK293T cells along with NF-κB firefly luciferase and control renilla luciferase reporter plasmids (see Materials and Methods), and NF-κB activity was measured at 18h post-transfection. Data were analyzed using one-way ANOVA with Tukey's multiple comparisons test. ** = p<0.01, **** = p<0.0001. Representative of two independent experiments with n=5-6 replicates per group.

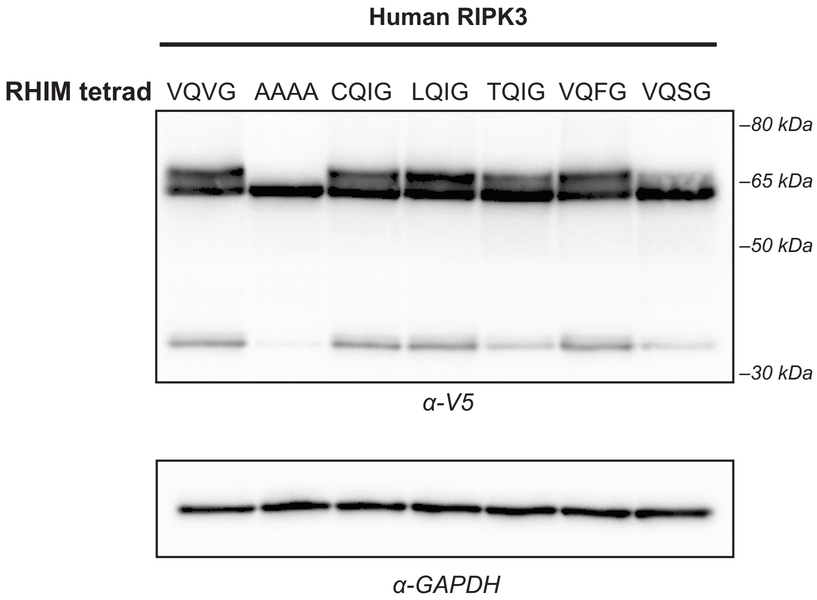


**Supplementary figure 15.** Expression of human RIPK3 RHIM variants. WT HEK293T cells were transfected with the indicated V5-RIPK3 protein. Protein expression was analyzed at 18 hours post-transfection by western blot using the indicated antibodies.

**Supplementary table 1.** Presence of ZBP1 and CASP8 in reptiles.

| Reptile | | ZBP1 | CASP8 |
|---|---|---|---|
| Crocodilians (Crocodilia) | **American alligator *(Alligator mississippiensis)*** | Yes | Yes |
| | **Chinese alligator *(Alligator sinensis)*** | Yes | Yes |
| | **Saltwater crocodile *(Crocadylus porosus)*** | Yes | Yes |
| | **Gharial *(Gavialis gangeticus)*** | Yes | Yes |
| Turtles & kin (Testudines) | **Loggerhead sea turtle *(Caretta caretta)*** | Yes | Yes |
| | **Green sea turtle *(Chelonia mydas)*** | Yes | Yes |
| | **Pinta island tortoise *(Chelonoidis abingdonii)*** | Yes | Yes |
| | **Painted turtle *(Chrysemys picta bellii)*** | Yes | Yes |
| | **Leatherback sea turtle *(Dermochelys coriacea)*** | Yes | Yes |
| | **Goode's thornscrub tortoise *(Gopherus evgoodei)*** | Yes | Yes |
| | **Bolson tortoise *(Gopherus flavomarginatus)*** | Yes | Yes |
| | **Yellow pond turtle *(Mauremys mutica)*** | Yes | Yes |
| | **Chinese pond turtle *(Mauremys reevesii)*** | Yes | Yes |
| | **Chinese softshell turtle *(Pelodiscus sinensis)*** | Yes | Yes |
| | **Three-toed box turtle *(Terrapene carolina triunguis)*** | Yes | Yes |
| | **Red eared slider *(Trachemys scripta elegans)*** | Yes | Yes |
| Snakes & lizards (Squamata) | Green anole *(Anolis carolinensis)* | No | No |
| | Tiger rattlesnake *(Crotalus tigris)* | No | No |
| | Schlegel's Japanese gecko *(Gekko japonicus)* | No | No |
| | Sand lizard *(Lacerta agilis)* | No | No |
| | Tiger snake *(Notechis scutatus)* | No | No |
| | Corn snake *(Pantherophis guttatus)* | No | No |
| | Common wall lizard *(Podarcis muralis)* | No | No |
| | Central bearded dragon *(Pogona vitticeps)* | No | No |
| | Brown-spotted pit viper *(Protobothrops mucrosquamatus)* | No | No |
| | Eastern brown snake *(Pseudonaja textilis)* | No | No |
| | Burmese python *(Python bivittatus)* | No | No |
| | Eastern fence lizard *(Scleroporus undulatus)* | No | No |
| | Townsend's least gecko *(Sphaerodactylus townsendi)* | No | No |
| | Western terrestrial garter snake *(Thamnophis elegans)* | No | No |
| | Common garter snake *(Thamnophis sirtalis)* | No | No |
| | Komodo dragon *(Varanus komodoensis)* | No | No |
| | Common lizard *(Zootoca vivipara)* | No | No |

**Supplementary Table 2.** Naturally-occurring RIPK3 RHIM tetrad variants. RIPK3 RHIM tetrad variants that deviate from the canonical motif (V/I-Q-V/I-G) at one or two residues were identified. Highlighted tetrads indicate those tested in Figure 4.

| Changes | Variant | # of species | Classification(s) |
|---|---|---|---|
| Single residue | TQIG | 1 | Lamprey |
| | LQIG | 5 | Fish, amphibian |
| | VQSG | 4 | Fish, amphibian |
| | CQIG | 1 | Fish |
| | VQFG | 7 | Fish, reptile, rodent, bat, eulipotyphlan |
| | IQYG | 3 | Fish |
| | VQCG | 1 | Fish |
| | FQIG | 2 | Amphibian |
| | VQLG | 2 | Fish |
| | IQFG | 1 | Bat |
| | IQLG | 5 | Ungulate, bat |
| | LQVG | 4 | Rodent, fish, |
| Two residues | CQFG | 1 | Amphibian |
| | FQRG | 1 | Fish |
| | FQNG | 3 | Fish |
| | FQCG | 1 | Fish |
| | FQKG | 1 | Fish |
| | LQSG | 2 | Fish |
| | LQLG | 9 | Fish |
| | FQLG | 1 | Fish |
| | LQTG | 1 | Fish |
| | FQSG | 7 | Fish |
| | AQFG | 2 | Fish |
| | FQHG | 8 | Fish |
| | LQQG | 3 | Fish |
| | LQHG | 4 | Fish |
| | FQYG | 4 | Fish |
| | FQQG | 5 | Fish |
| | LQCG | 1 | Fish |
| | LQFG | 13 | Fish |
| | LQYG | 6 | Fish |
